# A *Tm4sf1*-Marked Subpopulation of Endothelial Stem/Progenitor Cells Identified by Lung Single-Cell Omics of Pulmonary Arterial Hypertension

**DOI:** 10.1101/2022.01.09.475566

**Authors:** Jason Hong, Brenda Wong, Caroline Huynh, Brian Tang, Soban Umar, Xia Yang, Mansoureh Eghbali

## Abstract

**Rationale:** The identification and role of endothelial progenitor cells (EPCs) in pulmonary arterial hypertension (PAH) remains controversial. Single-cell omics analysis can shed light on EPCs and their potential contribution to PAH pathobiology.

**Objectives:** We aim to identify EPCs in rat lungs and assess their relevance to preclinical and human PAH.

**Methods:** Differential expression, gene set enrichment, cell-cell communication, and trajectory reconstruction analyses were performed on lung endothelial cells from single-cell RNA-seq of Sugen-hypoxia, monocrotaline, and control rats. Relevance to human PAH was assessed in multiple independent blood and lung transcriptomic datasets.

**Measurements and Main Results:** A subpopulation of endothelial cells (EA2) marked by *Tm4sf1*, a gene strongly implicated in cancer, harbored a distinct transcriptomic signature including *Bmpr2* downregulation that was enriched for pathways such as inflammation and angiogenesis. Cell-to-cell communication networks specific to EA2 were activated in PAH such as CXCL12 signaling. Trajectory analysis demonstrated EA2 has a stem/progenitor cell phenotype. Analysis of independent datasets revealed *Tm4sf1* is a marker for hematopoietic stem cells and is upregulated in PAH peripheral blood, particularly in patients with worse WHO functional class. EA2 signature genes including *Procr* and *Sulf1* were found to be differentially regulated in the lungs of PAH patients and in PAH models *in vitro*, such as BMPR2 knockdown.

**Conclusions:** Our study uncovered a novel *Tm4sf1*-marked stem/progenitor subpopulation of rat lung endothelial cells and demonstrated its relevance to preclinical and human PAH. Future experimental studies are warranted to further elucidate the pathogenic role and therapeutic potential of targeting EA2 and *Tm4sf1* in PAH.

## Introduction

Despite advances in our understanding of pulmonary arterial hypertension (PAH), it remains an incurable disease characterized by pathological pulmonary arterial remodeling and endothelial dysfunction. Endothelial progenitor cells (EPCs) are involved in endothelial homeostasis and angiogenesis and have been extensively studied in PAH animal models (1–3) and patients (4–10) using heterogenous approaches for EPC isolation by flow cytometry and/or culture. However, results have been conflicting and potentially confounding as to whether EPCs in PAH are protective vs. harmful (11), a reflection of the broader controversy in the stem cell field as to the precise definition and reliable isolation of EPCs (12). For example, conventional markers used to identify putative EPCs in numerous papers including PAH studies are now believed to be unreliable in isolating pure EPCs (12) and a specific marker for EPCs has yet to be identified (13).

The advent of single-cell omics can shed light on EPCs and their potential role in disease in a data-driven and unbiased approach. But while single-cell studies in PAH have recently been published including our recent work using lungs from PAH rat models (14–17), stem/progenitor endothelial cells have yet to be identified. However, rapidly advancing computational methods to dissect single-cell data provide an opportunity to unravel important biological discoveries. In this study of rat lung endothelial cells from single-cell RNA-sequencing (scRNA-seq), we uncover a *Tm4sf1*-marked subpopulation of endothelial stem/progenitor cells and demonstrate its relevance to PAH through its distinct transcriptomic signature, cell-to-cell signaling network, and differential regulation of its marker genes in circulating blood and lungs of PAH patients and animal models.

## Methods

Main methods are below with additional details provided in a supplement.

### Animals

Adult male Sprague-Dawley rats (250g-350g) were used for animal experiments, which were approved by the UCLA Animal Research Committee. As previously described (17), Sugen-hypoxia (SuHx) rats were injected subcutaneously with Sugen 5416 (20 mg/kg) followed by hypoxia at 10% O_2_ for 21 days and then normoxia for 14 days. Monocrotaline (MCT) rats were injected subcutaneously with MCT (60 mg/kg) followed by normoxia for 28 days. Age-matched control rats were kept in normoxia for 28 days. Lungs were then harvested and enzymatically dissociated into single-cell suspensions followed by scRNA-seq (17, 18) (*n* = 6/group) or snap-frozen for RNA isolation and bulk RNA-seq (*n* = 4/group).

### Differential expression analysis

Expression data was normalized, filtered, and clustered using Seurat R package (13)(19). Cell types were identified based on known cell type marker genes and endothelial clusters were then used for all downstream analyses. Differentially expressed genes (DEGs) were determined using MAST (20). To annotate DEGs, Gene Set Enrichment Analysis (GSEA) was performed using R package fgsea v1.18.0. PAH genes were retrieved from DisGeNET (21) and Comparative Toxicogenomics Database (CTD) (22).

### Cell-cell communication analysis

CellPhoneDB, a repository of ligands and receptors, was used to infer cell–cell communication from combined expression of multi-subunit ligand–receptor complexes in our scRNA-seq data (23). Enriched ligand–receptor interactions between two cell clusters are derived on the basis of expression of a receptor by one cell cluster and a ligand by another cell cluster. For each gene in the cluster, the percentage of cells expressing the gene and the gene expression mean are calculated. The expression levels of ligands and receptors within each cell cluster are considered and random shuffling is used to determine which ligand–receptor pairs display significant specificity between two cell clusters.

### Trajectory reconstruction

To predict differentiation states, we used CytoTRACE, a computational framework that leverages single-cell gene counts as a determinant of developmental potential, covariant gene expression, and local neighborhoods of transcriptionally similar cells to predict ordered differentiation states from scRNA-seq data (24).

### Independent datasets for validation

Select genes were queried in independent bulk and single-cell transcriptomic datasets of human lung and blood samples from Gene Expression Omnibus (GEO) or hosted on lab web servers.

## Results

### Tm4sf1-marked subpopulation of endothelial cells expresses a distinct transcriptomic signature relevant to PAH

The transcriptomes of 758 endothelial cells generated by scRNA-seq of 18 lungs from MCT, SuHx, and control rats (6/group) were clustered while retaining cell type annotations as previously described (17) yielding 3 distinct subclusters: 454 endothelial arterial type 1 cells (EA1), 255 endothelial arterial type 2 cells (EA2), and 49 endothelial capillary cells (Figure 1A). Each of the 3 subpopulations were represented in MCT, SuHx, and control lungs (Figure 1B). Subcluster-specific markers were identified such as *Nostrin* for EA1, *Tm4sf1* for EA2, and *Car4*, a known marker for EC (Figures 1C-1E). Arterial markers such as *Sox17* (Figure 1F) and *Efnb2* (Figure E1) were also identified confirming the arterial origin of EA1 and EA2 (25).

**Figure 1:**
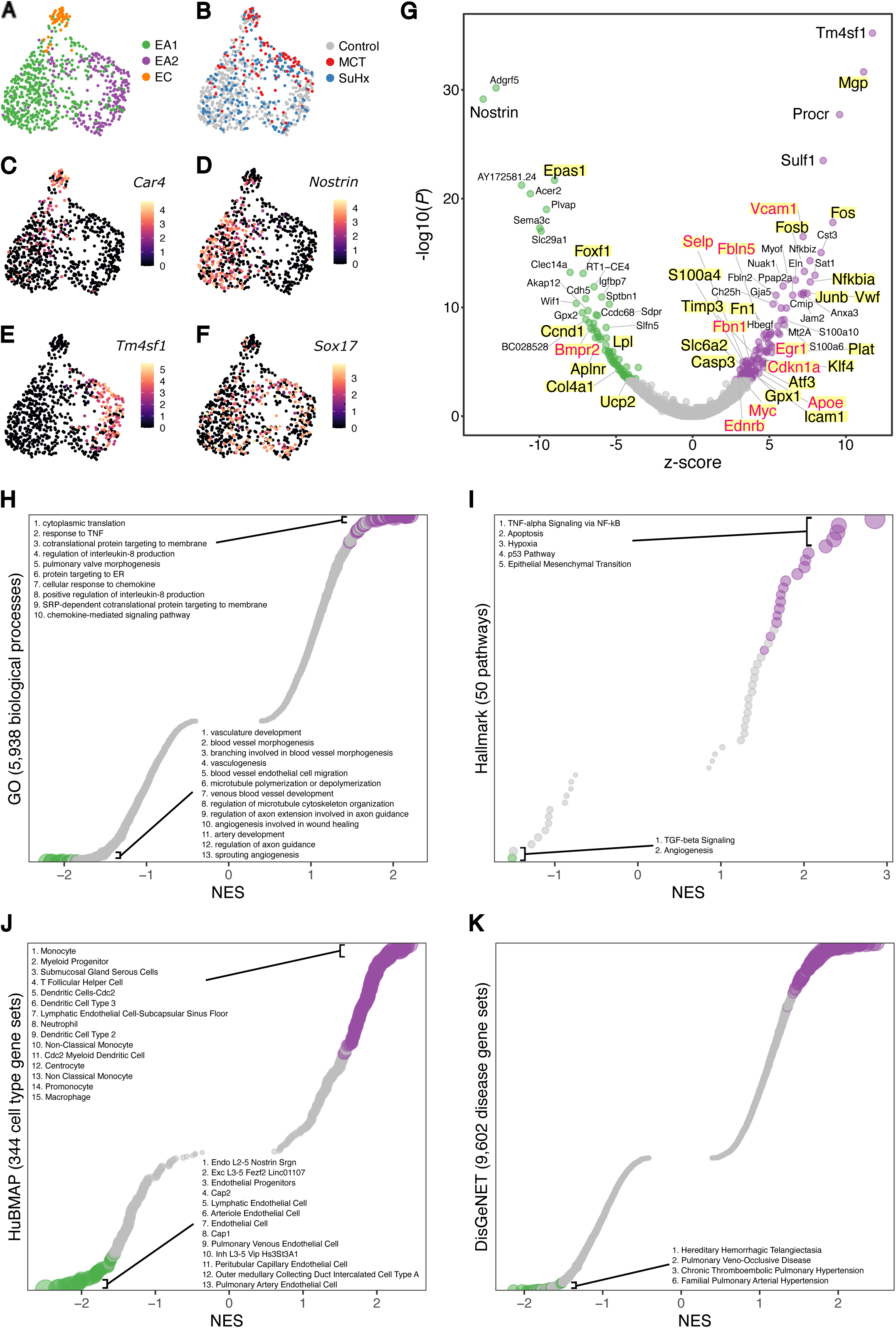
*Tm4sf1*-marked subpopulation of endothelial cells expresses a distinct transcriptomic signature relevant to PAH. (A-F) Uniform manifold approximation and projection (UMAP) plots showing clustering of 758 endothelial cell transcriptomes from 18 rats lungs with individual cells colored by (A) subpopulation, (B) disease condition (*n* = 6/group), and expression levels of the following marker genes: (C) *Car4* for endothelial capillary (EC), (D) *Nostrin* for endothelial arterial type 1 (EA), (E) *Tm4sf1* for endothelial arterial type 2 (EA2), and (F) *Sox17* as a known arterial marker expressed in both EA1 and EA2. Expression levels are represented as log normalized counts. (G) Volcano plot showing DEGs in 255 EA2 cells compared to 454 EA1 cells where x-axis represents MAST z-scores and y-axis indicates - log_10_(*P*). Significant upregulated (z > 0) or downregulated (z < 0) genes with FDR < 0.05 are shown as purple and green dots, respectively. Select top DEGs are labeled with their gene names. DEGs highlighted in yellow represent human PAH-associated genes from either (black text) or both (red text) CTD and DisGeNET databases. (H-K) Dots plots showing Gene Set Enrichment Analysis (GSEA) of the EA2 DEG signature using (H) GO, (I) Hallmark, (J) HuBMAP, and (K) DisGeNET gene sets where x-axis represents normalized enrichment scores (NES) in which NES greater than or less than zero and FDR < 0.05 represent gene sets significantly enriched in up- or downregulated genes, respectively and are colored in purple or green, respectively. The y-axis represents gene sets ordered by their NES. Select gene sets are labeled and numbered by their ordering as top gene sets enriched in up- or downregulated genes. Dots larger in size represent higher -log_10_(*FDR*) values. FDR = false discovery rate.

Given that the pathological hallmark of vascular remodeling in PAH predominantly occurs in the pulmonary arteries, we then evaluated the differences in gene expression between the two arterial subclusters EA1 and EA2. We found a total of 196 genes differentially expressed (DEGs; FDR < 0.05) with 117 genes upregulated and 79 genes downregulated in EA2 compared to EA1 (Figure 1G and Table E1). *Tm4sf1* was the top upregulated DEG in EA2 vs. EA1 and its expression was specific to EA2 when comparing all lung cell types (Figure E2). *Tm4sf1* encodes the cell-surface protein transmembrane 4 L six family member 1 which plays a role in the regulation of cell development, growth and motility and has been strongly implicated in various cancers (26) but has never before been studied in PAH. Given the importance of cancer-related processes and their overlap with PAH (27), we paid closer attention to the *Tm4sf1*-marked subpopulation of endothelial cells, EA2. *Nostrin* was the top downregulated gene in EA2 vs. EA1-it encodes nitric oxide synthase trafficking, an adaptor protein that binds to endothelial nitric oxide synthase (eNOS) and attenuates NO production. When intersecting other DEGs with disease-gene databases DisGeNET and CTD, we found that a number of DEGs are also known PAH genes such as *Bmpr2* which was downregulated in EA2 (Figure 1G).

We then used gene set enrichment analysis to determine what biological processes, pathways, cell types, and diseases are enriched in the gene signature of EA2 (vs. EA1). Among 5,938 biological processes from Gene Ontology (GO) (28), cytoplasmic translation and response to tumor necrosis factor (TNF) were most enriched in upregulated EA2 genes. Furthermore, angiogenesis-related processes were most enriched in downregulated EA2 genes (Figure 1H). Of the 50 Hallmark pathways (29), many known to be involved in PAH were enriched in genes upregulated in EA2 such as TNF*α*/NF-κB signaling, apoptosis, hypoxia, p53 pathway, and epithelial mesenchymal transition (EMT) (Figure 1I). To determine what cell type signatures may be enriched in EA2 vs. EA1 DEGs, we tested 344 cell type-specific gene sets from HubMAP (30) and unexpectedly found that immune cell signatures, particularly from monocytes, were most enriched in genes differentially upregulated in EA2 whereas endothelial signatures were most enriched in genes upregulated in EA1 (Figure 1J). Additionally, we found that across 9,602 disease-specific gene sets from DisGeNET, genes downregulated in EA2 vs. EA1 were most enriched for diseases associated with primary or secondary forms of pulmonary hypertension (PH): hereditary hemorrhagic telangiectasia (HHT), pulmonary veno-occlusive disease (PVOD), chronic thromboembolic pulmonary hypertension (CTEPH), and familial PAH (Figure 1K). Thus, not only was *Bmpr2* downregulated in EA2 as it is in human PAH lungs (31), the downregulated portion of the EA2 signature on a transcriptome-wide scale was also highly associated with PAH.

### EA2-specific cell-to-cell signaling networks are activated in PAH

Having identified a transcriptionally distinct subpopulation of lung endothelial cells relevant to PAH, we next hypothesized that communication between EA2 and other cell types occurs in a disease-specific manner given that many other cell types besides endothelial cells have also been implicated in the complex pathobiology of PAH. Using CellPhoneDB to predict ligand-receptor interactions between cell types, we found overall increased intercellular communication between various immune, mesenchymal, and epithelial cells in the lungs of PAH rat models compared to control (Figure 2A). When quantifying the number of statistically significant ligand-receptor interactions between endothelial cells and other cell types, EA2 exhibited stronger cell-to-cell signaling in MCT and SuHx lungs compared to control that was not seen in EA1 nor EC (Figure 2B).

**Figure 2:**
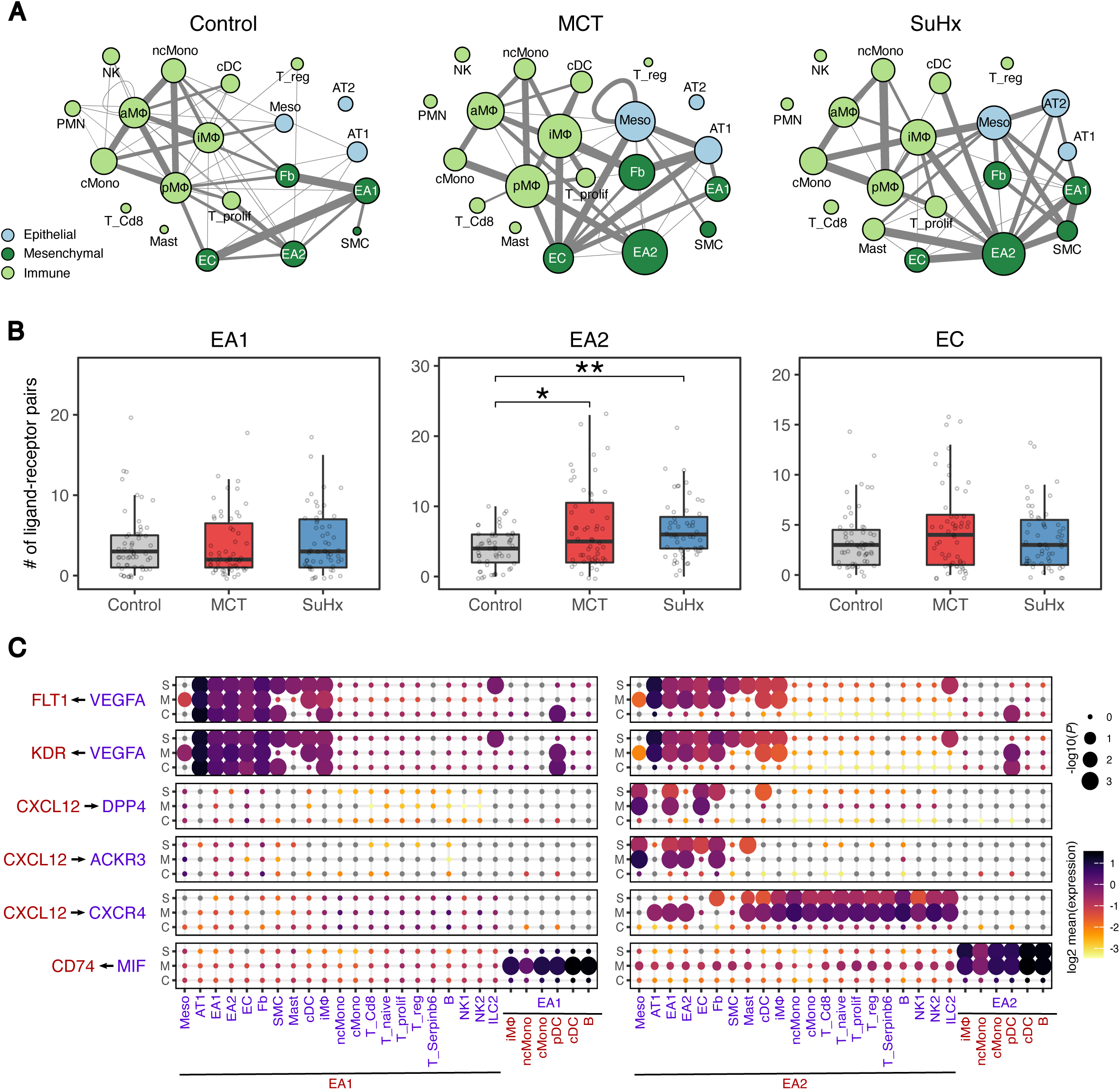
EA2-specific cell-to-cell signaling networks are activated in PAH. (A) Network visualizations of control, MCT, and SuHx lungs where nodes are represented as cell types and edge weight is proportional to the number of distinct and statistically significant ligand-receptor interaction pairs between two cell types as determined by CellPhoneDB. Thicker edges correspond to a higher number of ligand-receptor pairs. Node size is proportional to the sum of edge weights representing a given cell type’s interactions with all other cell types. The top 10% of edges by weight are shown for each condition. (B) Box plots comparing cell-cell interactions where each dot represents the number of statistically significant ligand-receptor interaction pairs between endothelial subpopulations (EA1, EA2, and EC) and all other cell types in control, MCT, and SuHx lungs. Wilcoxon rank-sum test: \**P* < 0.05 and \*\**P* < 0.01. (C) Dot plots showing the log2 mean expression of select ligand-receptor pairs in control (C), MCT (M), and SuHx (S) between select cell types and EA1 or EA2. Size of dots are proportional to strength of p-value. aMΦ = alveolar macrophages; cMono = classical monocytes; cDC = conventional dendritic cells; Fb = fibroblasts; ILC2 = type 2 innate lymphoid cells; iMΦ = interstitial macrophages; meso = mesothelial, ncMono = non-classical monocytes; NK1 = NK cells 1; NK2 = NK cells 2; PMN = neutrophils; pDC = plasmacytoid dendritic; pMΦ = proliferating macrophages; SMC = smooth muscle cells; T_prolif = proliferating T cells; T_reg = regulatory T cells.

We further examined specific ligand-receptor interactions and found that signaling molecules known to play a role in PAH such as vascular endothelial growth factor A (VEGFA) (32), C-X-C motif chemokine ligand 12 (CXCL12) (33, 34), and macrophage migration inhibitory factor (MIF) (35, 36) were not only differentially active in PAH models compared to control but also specific to cell-cell communication involving EA2 but not EA1 (Figure 2C) nor EC (Figure E3). For instance, VEGFA from cell types such as fibroblasts and interstitial macrophages were predicted to interact in a PAH- and EA2-specific manner with its cognate receptors fms related receptor tyrosine kinase 1 (FLT1; VEGFR-1) and kinase insert domain receptor (KDR; VEGFR-2). CXCL12 signaling was highly specific to EA2 with distinct cell-cell communication profiles specific to the particular receptor expressed on target cell types. For example, EA2’s CXCL12 interaction with C-X-C motif chemokine receptor 4 (CXCR4) was observed in various PAH immune cell types of both lymphoid and myeloid lineage whereas its interaction with atypical chemokine receptor 3 (ACKR3; CXCR7) and dipeptidyl peptidase 4 (DPP4) was noted in PAH fibroblasts, endothelial, and mesothelial cells (Figure 2C). MIF is a key pro-inflammatory immune regulator whose interaction with its receptor CD74 on myeloid and B cells was statistically significant for both MCT and SuHx EA2 compared to control, a pattern that was unique to EA2 (Figure 2C).

### Trajectory analysis reveals that EA2 has a stem/progenitor cell phenotype

Given the important role of CXCL12 and its chemokine receptor CXCR4 in the homing and function of endothelial progenitor cells (EPCs) (9, 37–40) and the specificity of this signaling axis for EA2 communication in our rat lung scRNA-seq analysis, we asked whether EA2 might have a stem/progenitor cell phenotype. We used CytoTRACE (24), an unbiased computational method that leverages the number of expressed genes per cell as a determinant of developmental potential to predict the differentiation state of single cells from our endothelial scRNA-seq data (Figure 3A). Compared to EA1 and EC cells, we found that EA2 cells had significantly higher CytoTRACE scores corresponding to a less differentiated state (Figure 3B). Comparing CytoTRACE scores by condition, endothelial cells from MCT and SuHx lungs were predicted to be less differentiated compared to control, consistent with prior evidence implicating endothelial progenitor cells in PAH (9, 41, 42). To identify markers of the less differentiated phenotype, all 12,116 genes with detectable expression in our dataset were rank-ordered on the basis of their correlation with CytoTRACE scores across all 758 endothelial cells analyzed. Ribosomal genes correlated most strongly with a less differentiated state (Figure 3D) and thus were downregulated with a more differentiated state, consistent with prior studies (43, 44). Gene set enrichment analysis revealed many GO biological processes significantly enriched in genes correlated with a less differentiated state compared to a more differentiated state (219 vs. 0 out of 5,994 total gene sets) (Figure E4). Top enriched biological processes included cotranslational protein targeting to membrane and cytoplasmic translation which were also top enriched GO biological processes of the EA2 gene signature (Figure 1H) providing further evidence for a stem/progenitor phenotype of EA2.

**Figure 3:**
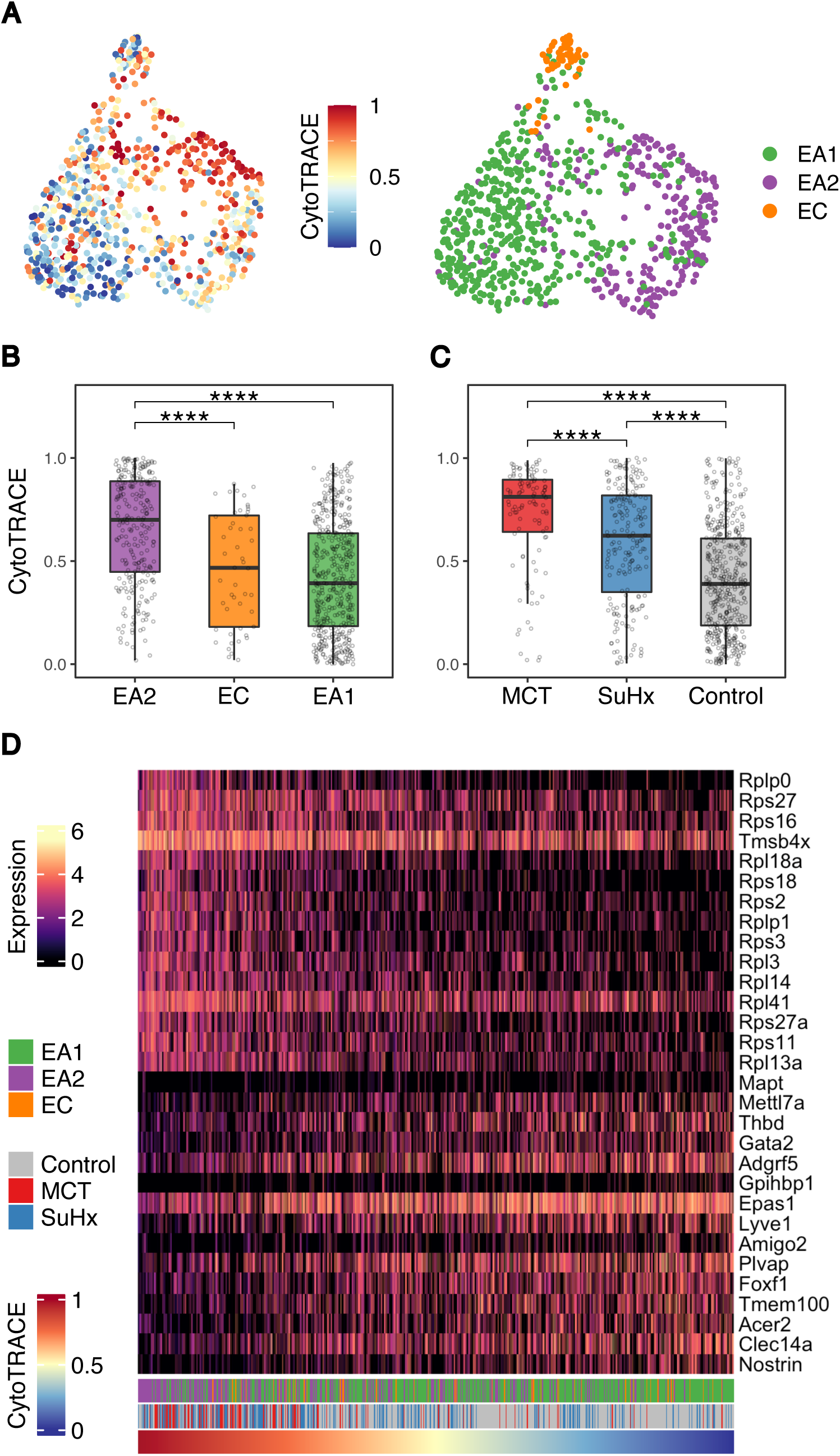
Trajectory analysis reveals that EA2 has a stem/progenitor cell phenotype. (A) UMAP plots showing clustering of endothelial cells colored by CytoTRACE score (left) and cell type (right). Higher CytoTRACE scores represent a less differentiated state. (B-C) Box plots comparing cell CytoTRACE scores grouped by (B) endothelial subpopulation and (C) disease condition. Wilcoxon rank-sum test: \*\*\*\**P* < 0.0001. (D) Heatmap showing normalized expression of top genes most positively or negatively correlated with cell CytoTRACE scores. Each column is a cell ordered from left to right by decreasing CytoTRACE scores. Cell annotations for cell type and condition are shown at the bottom.

### Tm4sf1 is a marker for hematopoietic stem cells and is upregulated in the peripheral blood of PAH patients

Given prior studies implicating circulating EPCs in PAH patients (6–9), we asked whether *Tm4sf1*, the most highly specific marker for rat lung EA2, could also be a marker for circulating stem/progenitor cells in human PAH. Using a dataset of 7,551 human blood cell transcriptomes representing 43 cell type clusters derived from 21 healthy donors (45), we found that *TM4SF1* was specifically expressed in and one of 19 marker genes for hematopoietic stem cell/multipotent progenitor 1 cells (HSC/MPP1) (FDR = 2.0e-15) (Figure 4A) further suggesting that *Tm4sf1*-marked EA2 cells may be a stem/progenitor subpopulation of endothelial cells.

**Figure 4:**
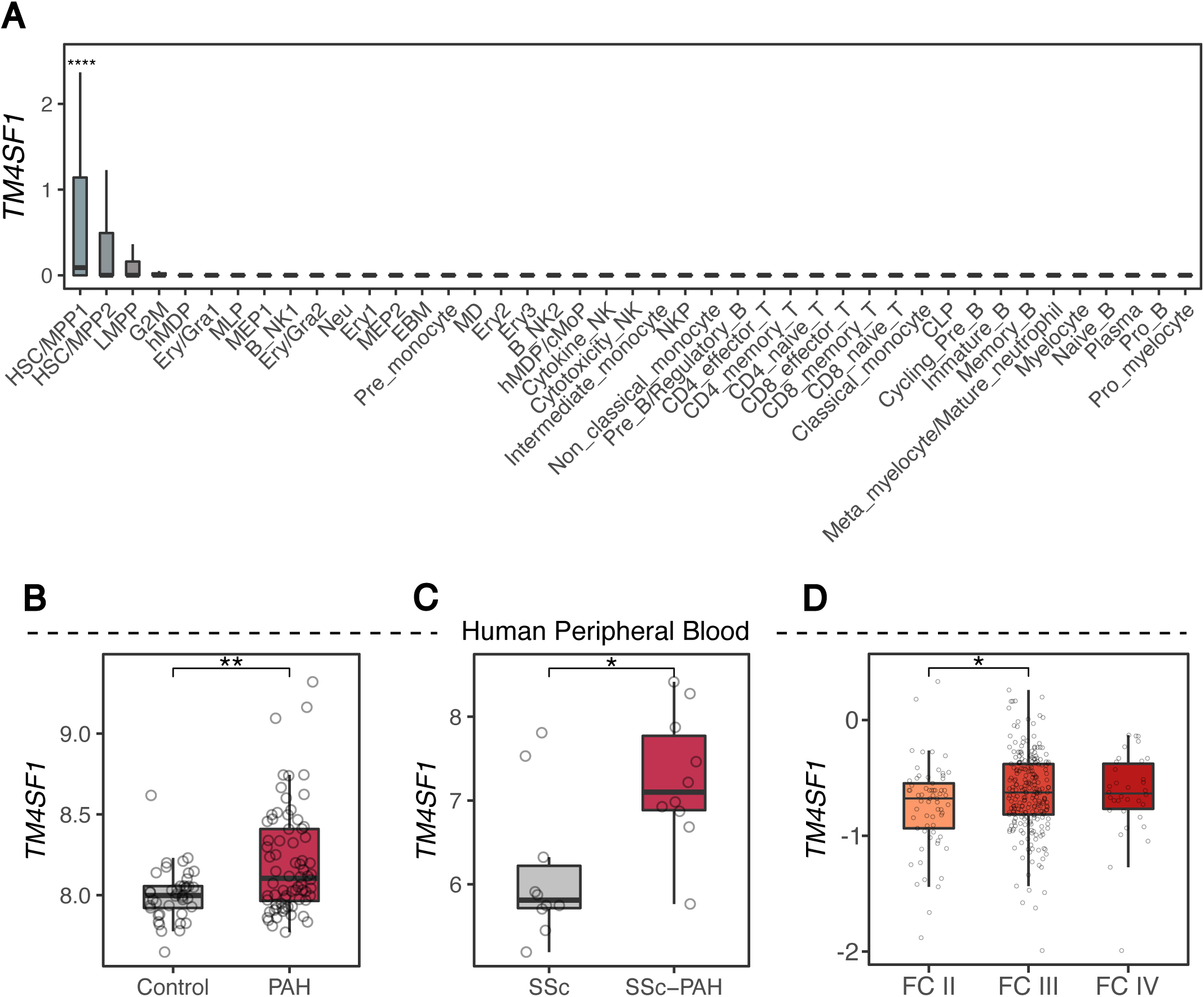
*Tm4sf1* is a marker for hematopoietic stem cells and is upregulated in the peripheral blood of PAH patients. (A) Box plots showing the expression of *TM4SF1* in a dataset of 7,551 human blood cell transcriptomes representing 43 cell type clusters derived from 21 healthy donors (45). (B-D) Box plots showing expression of *TM4SF1* in PBMCs of (B) 72 PAH patients compared to 41 healthy control (46) and (C) 10 SSc-associated PAH patients compared to 10 SSc without PAH (47) and (D) in whole blood of 65 FC II, 239 FC III, and 34 FC IV PAH patients. *P* values in (B) and (C) were obtained from GEO2R and determined by Wilcoxon rank-sum test in (D): \**P* < 0.05 and \*\**P* < 0.01.

Supporting the relevance of this marker to PAH, we found in two independent microarray datasets that *TM4SF1* was significantly upregulated in circulating peripheral blood mononuclear cells (PBMCs) of PAH patients compared to healthy control (*n* = 72 and 41, respectively) (46) and in systemic sclerosis (SSc)-associated PAH patients compared to SSc without PAH (*n* = 10 and 10, respectively) (47) (Figures 4B-4C). Furthermore, in a third dataset of whole blood RNA-seq, PAH patients with higher *TM4SF1* expression were associated with worse WHO functional class (FC) (*n* = 65, 239, and 34 for FC II, III, and IV, respectively) (Figure 4D).

### EA2 signature genes are differentially regulated in the lungs of PAH patients and in PAH models of human endothelial cells

Having established that the EA2 marker *Tm4sf1* is upregulated in PAH peripheral blood and associated with disease severity, we next investigated whether *Tm4sf1* and other EA2 markers are dysregulated in PAH lungs. Given that the power to detect PAH-specific differences in *Tm4sf1* expression in our rat lung scRNA-seq dataset may have been limited by relatively lower yield of endothelial cells from the tissue dissociation into single cells, we evaluated whether *Tm4sf1* expression is dysregulated at the bulk lung tissue level. We performed bulk RNA sequencing on lung tissue from a subset of the same animals used for scRNA-seq (n = 4/group) and found that *Tm4sf1* was significantly upregulated in the lungs of both MCT and SuHx compared to control (Figure 5A). Supporting the human relevance of this finding, a 2006 study from Harefield Hospital in the United Kingdom which used suppression subtractive hybridization found that *TM4SF1* was one of 27 upregulated genes in PAH vs. donor lungs (*n* = 2/group) (48). However, we probed a 2010 University of Pittsburgh (Pitt) microarray dataset (49) and found that *TM4SF1* was instead downregulated in PAH lungs compared to control (*n* = 18 and 13, respectively) (Figure 5B) which could reflect differences in patient population and disease severity between datasets. Furthermore, analysis of a 2020 Pitt scRNA-seq dataset revealed that *TM4SF1* was downregulated in endothelial cells from PAH lungs compared to donors (*n* = 3 and 6, respectively) (15) (Figure 5C). We then evaluated two other top EA2 markers *Procr* and *Sulf1* and discovered both to be upregulated in the lungs of PAH patients in two independent microarray datasets each (Figures 5D-5G) (49–51).

**Figure 5:**
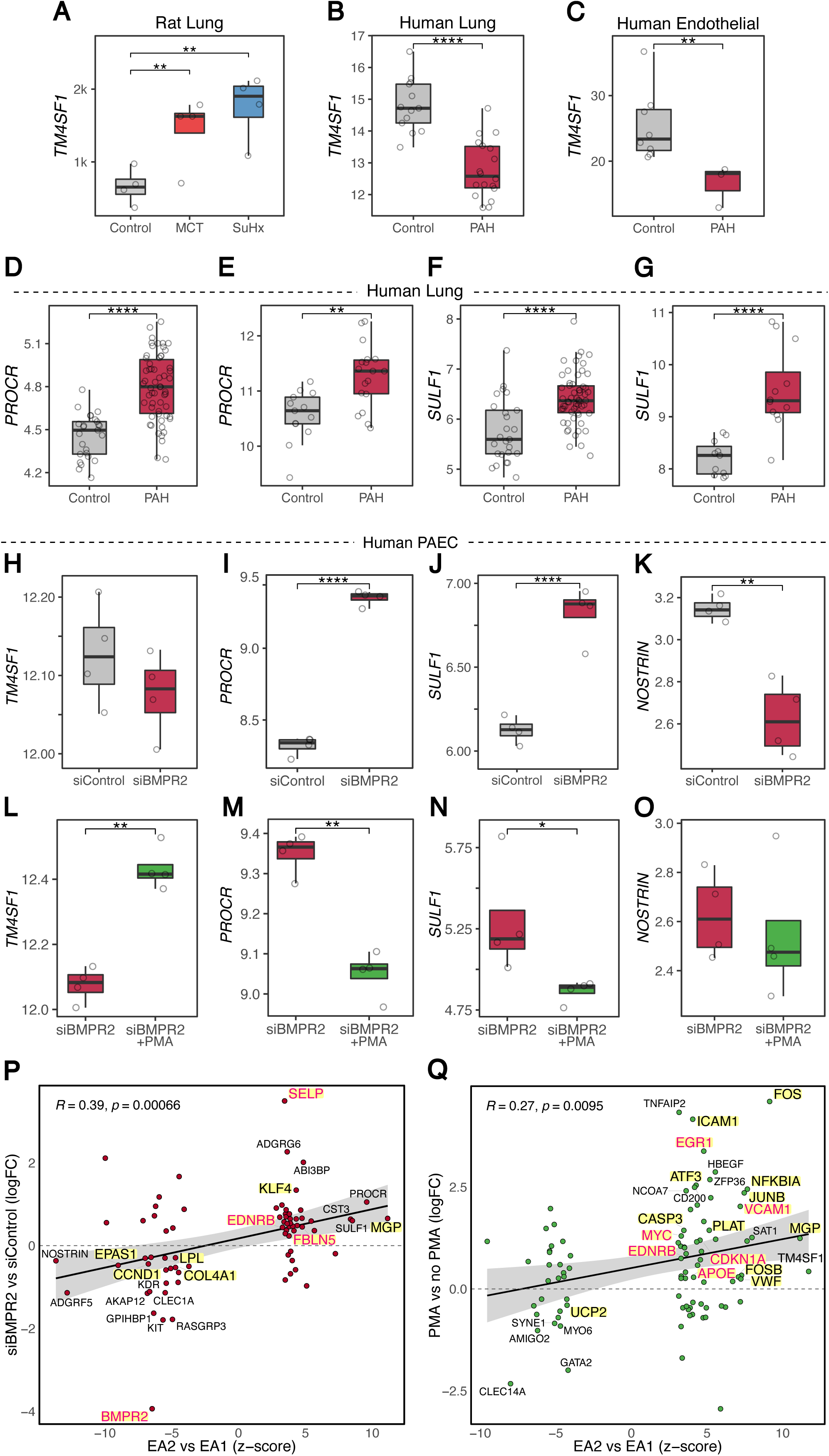
EA2 signature genes are differentially regulated in the lungs of PAH patients and by PAH models in human endothelial cells. (A-C) Box plots showing *TM4SF1* expression in (A) control, MCT, and SuHx lungs (*n* = 4/group), in (B) 18 PAH patient lungs vs 13 controls (49), and in (C) averaged endothelial scRNA-seq cells from 6 PAH patient lungs vs 3 controls (15). (D, F) Box plots showing expression of EA2 markers *PROCR* and *SULF1* in 58 PAH patient lungs vs 25 controls (50). (E) Box plot showing expression of *PROCR* in 18 PAH patient lungs vs 13 controls (49). (G) Box plot showing expression of *SULF1* in 12 PAH patient lungs vs 11 controls (51). (H-K) Box plots showing expression of (H) *TM4SF1*, (I) *PROCR*, (J) *SULF1*, (K) *NOSTRIN* in 4 *BMPR2*- vs 4 control-siRNA-transfected primary human PAECs from 4 donors (52). (L-O) Box plots showing expression of (L) *TM4SF1*, (M) *PROCR*, (N) *SULF1*, (O) *NOSTRIN* in 4 *PMA*-exposed vs 4 nonexposed *BMPR2*-silenced PAECs from 4 donors (52). *P* values were obtained from GEO2R for (B, D-O), determined by DESeq2 for (A) and by Wilcoxon rank-sum test for (C): \**P* < 0.05, \*\**P* < 0.01, \*\*\**P* < 0.001, ****FDR < 0.05. (P-Q) Scatter plots showing correlation between DEGs in EA2 vs. EA1 (x-axis) and DEGs from either (P) *BMPR2-*silenced PAECs (*n* = 8/group) or (Q) PMA-exposed PAECs (*n* = 8/group) (52) with select top DEGs labeled by their gene names. DEGs highlighted in yellow represent human PAH-associated genes from either (black text) or both (red text) CTD and DisGeNET databases. PMA = phorbol 12-myristate 13-acetate; logFC = log fold change; FDR = false discovery rate.

Having determined that top EA2 markers are dysregulated in the lungs of PAH patients and given that *Bmpr2*, the most well-established causal gene in PAH, is downregulated in EA2 compared to EA1 (Figure 1G), we next asked whether EA2 markers are regulated by *Bmpr2*. We analyzed a microarray of siRNA knockdown of *BMPR2* in primary human pulmonary artery endothelial cells (PAECs) *in vitro* (52) and found no change in *TM4SF1* but *PROCR* and *SULF1* were upregulated consistent with upregulation in EA2 vs. EA1 and PAH vs. donor lungs (Figures 5H-5K). Furthermore, *NOSTRIN*, the top downregulated gene in EA2 vs. EA1 (Figure 1G), was also downregulated in human PAECs upon *BMPR2* knockdown (Figure 5K).

Given the strong connection between *Tm4sf1* and various cancers (26), we then evaluated the response of *Tm4sf1* and other signature genes of EA2 to phorbol 12-myristate 13-acetate (PMA), a potent tumor promoter which activates protein kinase C and downstream mitogen-activated protein kinase (MAPK) pathways which have been implicated in PAH pathogenesis (52–54). We found that *TM4SF1* was upregulated in *BMPR2*-silenced human PAECs exposed to PMA (Figure 5L), while *PROCR*, *SULF1*, and *NOSTRIN* were either downregulated or unchanged (Figures 5M-5O). Overall, we found significant correlation between DEGs in EA2 (vs. EA1) and DEGs in human PAECs after either *BMPR2* knockdown (Figure 5P) or PMA stimulation (Figure 5Q) suggesting that portions of the EA2 signature can be explained by both downregulation of *Bmpr2* and stimulation with a tumor promoter. Overlapping DEGs were enriched for known PAH genes (Figures 5P-5Q) and pathways (Figures E6A-E6B) in both conditions, such as EMT in *BMPR2*-silenced PAECs and TNF*α*/NF-κB signaling in PMA-simulated PAECs.

## Discussion

In this study, we uncover a *Tm4sf1*-marked subpopulation of rat lung endothelial cells and demonstrate its relevance to PAH through its distinct transcriptomic signature, cell-to-cell signaling network, stem/progenitor cell phenotype, and differential regulation of its marker genes in circulating blood and lungs of PAH patients and animal models.

*Tm4sf1*, the top marker of EA2, encodes a cell-surface protein and member of the transmembrane 4 superfamily which mediates signal transduction events that regulate cell development, activation, growth, and motility(55). TM4SF1 is highly expressed and associated with poor survival in a wide range of cancers (26) but has never previously been studied in PAH, a disease that shares many pathobiological features with cancer for which many antineoplastic therapies have been evaluated for potential repurposing in PAH (27). With the accumulating evidence for TM4SF1’s role in cancer, potential oncologic therapies have been investigated such as in a phase 1/2 clinical trial evaluating chimeric antigen receptor T-cells (CAR-T) to treat patients with TM4SF1-positive pancreatic, colorectal, gastric, and lung cancer (ClinicalTrials NCT04151186). TM4SF1 promotes the migration and invasion of cancer cells by inducing EMT, self-renewal ability, and tumor angiogenesis (26), all of which are mechanisms relevant to the pathological pulmonary vascular remodeling in PAH.

Similar to malignant tumors, we found *Tm4sf1* to be upregulated in the lungs of SuHx and MCT rats (at the standard time points of 5 total weeks and 4 weeks, respectively) and in the lungs of PAH patients undergoing transplant from a 2006 UK study. However, *TM4SF1* was downregulated at the lung tissue level in a 2010 Pitt microarray and at the lung endothelial cell level in a 2020 Pitt scRNA-seq dataset in explanted PAH lungs. Differences in the directionality of *TM4SF1’s* dysregulation in PAH could reflect institutional, national, and/or temporal differences in the characteristics of patients undergoing lung transplant such as disease stage, severity, and/or treatment history. We hypothesize that TM4SF1 is upregulated earlier in the disease course and then becomes downregulated in end-stage PAH, both of which may be harmful. Supporting this hypothesis, knockdown of TM4SF1 in PAECs causes endothelial senescence (56) which has recently been shown to be causal in the transition from a reversible to the irreversible pulmonary vascular phenotype of end-stage PAH (57).

Furthermore, we found that not only was *TM4SF1* dysregulated in PAH lungs, it was upregulated in PAH peripheral blood in multiple independent cohorts, associated with disease severity, and a highly specific marker for HSCs, which reside in bone marrow, circulate in peripheral blood, and give rise to not only all blood cell lineages, but also EPCs (58). Given that *Tm4sf1* was also a highly specific marker for EA2 in our rat lung dataset, this raises the possibility that EA2 is a stem/progenitor subpopulation of lung endothelial cells that derives from bone marrow via the circulating blood. Consistent with a stem/progenitor phenotype, our trajectory analysis also predicted that EA2 cells are of an earlier differentiation state compared to other endothelial cells. Supporting this finding, a recent study identified *Tm4sf1* as a marker of multipotent endothelial precursor cells by scRNA-seq of lineage-traced mouse lungs (59).

While controversy and uncertainty persists in the definition and isolation of EPCs and their role in PAH (11–13, 60), it is generally accepted there exists 2 distinct subsets: so-called late EPCs derived from endothelial lineage and early EPCs derived from hematopoietic lineage with which EA2 shares features. Not only does EA2 share a novel marker with HSCs supporting its hematopoietic origin, we also unexpectedly found that the EA2 signature is highly enriched for myeloid signatures (Figure 1J) which is similarly noted in early EPCs (61). In fact, renaming of early EPCs to “myeloid angiogenic cells” has been proposed to more precisely define its phenotype and function (12) such as secretion of key proangiogenic factors like CXCL12 (60) - a chemokine implicated in PAH (33, 34, 62, 63) whose interaction with various target cells were unique to EA2 (Figure 2C). Consistent with our finding that EA2’s marker *TM4SF1* is upregulated in PAH circulation, a previous study found that early EPCs, but not late EPCs, are increased in PAH circulation as well (8). In other studies of early EPCs, one found a trend towards an increase in PAH (7), and the other found a decrease in PAH (6) but used an older isolation method now believed to be unreliable in EPC purification (13).

Further indicating EA2’s relevance in PAH pathobiology, we also demonstrated that two other top EA2 markers, *Sulf1* and *Procr*, were upregulated in explanted PAH lungs in multiple independent cohorts. In line with our findings, SULF1 (Sulfatase-1) was recently shown to be upregulated in remodeled pulmonary arterioles in PAH patients and MCT and SuHx rats (64). Further supporting EA2’s stem/progenitor phenotype, *Procr,* encoding protein C receptor, was previously identified as a marker for murine HSCs (65) and blood vascular endothelial stem cells (66) which displayed EMT and angiogenic signatures. While upregulated EA2 genes (relative to EA1) were enriched for EMT, genes downregulated in EA2 were enriched for angiogenesis (Figures 1H-1I). For example, the top downregulated gene in EA2, NOSTRIN (nitric oxide synthase trafficking), is known to inhibit angiogenesis (67) thus supporting an angiogenic phenotype of EA2. Furthermore, NOSTRIN has also been shown to suppress TNF*α*/NF-κB signaling (67) which is congruent with TNF*α*/NF-κB signaling as the top Hallmark pathway upregulated in the EA2 signature (Figure 1I).

Another downregulated gene in EA2 was *Bmpr2*, the most well-established causal PAH gene whose expression is also decreased in patient lungs of various PAH subtypes including non *BMPR2* mutation-related PAH (31). Reduced BMPR2 expression is known to cause exaggerated TGF-beta signaling in line with our finding that TGF-beta signaling was the top Hallmark pathway enriched in genes downregulated in EA2 (Figure 1I). Overactive TGF-beta signaling ultimately leads to pulmonary vascular remodeling in PAH through processes such as inflammation, angiogenesis, and EndMT (68), all of which are also upregulated in EA2. Furthermore, we demonstrated that *Sulf1*, *Procr*, *Nostrin* and a number of other genes are all differentially regulated after *BMPR2* knockdown in human PAECs in the same direction as EA2’s signature suggesting these genes may be downstream of BMPR2 signaling and thus downregulation of *Bmpr2* likely induces a portion of EA2’s gene signature. *TM4SF1* was unchanged upon *BMPR2* knockdown suggesting it may act upstream or independently of BMPR2 signaling, or a difference was not detected due to variability in experimental replicates. However, *TM4SF1* and many other EA2 signature genes including known PAH genes were upregulated upon PMA stimulation suggesting their potential regulation by MAPK signaling pathways which have been implicated in PAH pathogenesis (52–54).

Given the challenges of enzymatically dissociating tissue into single cells that may be particularly fragile or tightly embedded, we cannot exclude the possibility that other arterial endothelial populations were not captured in our scRNA-seq dataset. Alternative methods such as single-nucleus RNA-seq may provide a more precise cellular representation of tissue composition (69).

In conclusion, our study provides an in-depth analysis of rat lung endothelial heterogeneity using unbiased single-cell computational methods to uncover a novel *Tm4sf1*-marked stem/progenitor subpopulation with high relevance to PAH supported by multiple independent human and rat datasets. Future experimental studies are warranted to further elucidate the pathogenic role and therapeutic potential of targeting EA2 and *Tm4sf1* in PAH.

## Acknowledgements

The authors would like to express gratitude to Dr. Douglas Arneson for his guidance with scRNA-seq analytical techniques, Drs. In Sook Ahn and Graciel Diamante for their help in performing Drop-seq, Drs. Christine Cunningham and May Bhetraratana for their contribution to lung tissue dissociation, and Dr. Gregoire Ruffenach for his assistance with the animal experiments.

## Supplementary Methods

### Tissue processing

Lungs from control, SuHx and MCT rats were harvested after perfusion of lungs with gravity infusion of PBS via an 18g needle through the RV as previously described (1). The left lung was used for scRNA-seq and both lungs used for FACS. The tissue was minced with scissors prior to enzyme dissociation. For the first nine rats (*n* = 3/group), collagenase/dispase (Sigma, cat. no. 10269638001) at 1 mg/ml in PBS was used as previously described (2), with incubation at 37 °C for 45 min under continuous horizontal shaking (300 r.p.m.) For the second (*n* = 3/group for scRNA-seq) and third (*n* = 4/group for FACS) sets of rats, minced lungs were incubated with collagenase I (Worthington, cat. no. LS004194) at 1 mg/ml in DMEM + 10% FBS in a 37 °C water bath for 60 min, with trituration every 5 minutes (1–3)(3). The use of different cell dissociation protocols helps resolve potential biases in tissue dissociation and recover a more comprehensive atlas of lung cell types. After incubation, dissociated tissues were sequentially passed through 70- and 40-micron cell strainers with centrifugation at 300g for 5 min at 4 °C to pellet cells in between each step. Cells were then treated with RBC lysis buffer.

### Single-cell RNA-seq barcoding and library preparation

Lung single cells from SuHx, MCT, and control rats (*n* = 6/group) were suspended at a final concentration of 100 cells/microliter in 0.01% BSA-PBS as previously described (1). Barcoded single cells, or STAMPs (single-cell transcriptomes attached to microparticles), and cDNA libraries were generated following the Drop-seq protocol from Macosko et al. (4) and version 3.1 of the online Drop-seq protocol [http://mccarrolllab.org/download/905/]. Briefly, single cell suspensions at 100 cells/μl, EvaGreen droplet generation oil (BIO-RAD, Hercules, CA, USA), and ChemGenes barcoded microparticles (ChemGenes, Wilmington, MA, USA) were co-flowed through a FlowJEM aquapel-treated Drop-seq microfluidic device (FlowJEM, Toronto, Canada) at recommended flow speeds (oil: 15,000 μl/hr, cells: 4000 μl/hr, and beads 4000 μl/hr) to generate STAMPs. The following modifications were made to the online published protocol to obtain enough cDNA as quantified by TapeStation (Agilent) to continue the protocol: (1) the number of beads in a single PCR tube was 4000, (2) the number of PCR cycles was 4 + 11 cycles, and (3) multiple PCR tubes were pooled. The libraries were then checked on TapeStation for library quality, average size, and concentration estimation. The samples were then tagmented using the Nextera DNA Library Preparation kit (Illumina) and multiplex indices were added. After another round of PCR, the samples were checked on TapeStation for library quality check before sequencing. (A cell doublet rate of 5.6% was obtained by running the microfluidic device without the lysis buffer and counting the percentage of cell doublets through three separate runs.)

### Illumina high-throughput sequencing of single-cell RNA-seq libraries

The Drop-seq library molar concentration was quantified by Qubit Fluorometric Quantitation (Thermo Fisher) and library fragment length was estimated using TapeStation as previously described (1). Sequencing was performed on an Illumina HiSeq 4000 (Illumina) instrument using the Drop-seq custom read 1B primer (GCCTGTCCGCGGAAGCAGTGGTATCAACGCAGAGTAC) (IDT, Coralville, IA, USA). Paired end reads were 100bp in length. Read 1 consists of the 12 bp cell barcode, followed by the 8 bp UMI, and the last 80 bp on the read are not used. Read 2 contains the single cell transcripts.

### Single-cell RNA-seq data pre-processing and quality control

Demultiplexed fastq files generated from Drop-seq were processed to digital expression gene matrices using Drop-seq tools version 1.13 (https://github.com/broadinstitute/Drop-seq) as previously described (1). The workflow is available on github (https://github.com/darneson/dropSeqPipeDropEST). Briefly, fastq files were converted to BAM format and cell and molecular barcodes were tagged. Reads corresponding to low quality barcodes were removed and any occurrence of the SMART adapter sequence or polyA tails found in the reads was trimmed. These cleaned reads were converted back to fastq format to be aligned to the rat reference genome Rnor 6.0 using STAR-2.5.0c. After the reads were aligned, the reads which overlapped with exons, introns, and intergenic regions were tagged using a RefFlat annotation file of Rnor 6.0. The count values for each cell were normalized by the total number of UMIs in that cell and then multiplied by 10,000 and log transformed. Single cells were identified from background ambient mRNA using thresholds of at least 200 genes, 400 transcripts and no more than 50% mitochondrial fraction.

### Identification of cell clusters

The Seurat R package version 3.0.0.9000 (https://github.com/satijalab/seurat) was used to project all sequenced cells onto two dimensions using Uniform Manifold Approximation and Projection (UMAP) (5) and Louvain clustering (6) was used to assign clusters as previously described (1). The optimal number of principal components used for UMAP dimensionality reduction and Louvain clustering was determined using the Jackstraw permutation approach. The density used to assign clusters was identified using a parameter grid search.

### Identification of marker genes and cell types of individual cell clusters

We defined cell cluster specific marker genes from our Drop-seq dataset using the FindConservedMarkers function in Seurat across all the samples as previously described (1). Briefly, a Wilcoxon rank-sum test is run within each sample and a meta p-value across all samples is computed to assess the significance of each gene as a marker for a cluster. Within each sample, the cells are split into two groups: single cells from the cell type of interest and all other single cells. To be considered in the analysis, the gene had to be expressed in at least 10% of the single cells from one of the groups and there had to be at least a 0.25 log fold change in gene expression between the groups. This process was conducted within each sample separately, and then a meta p-value was assessed from the p-values across all samples. Multiple testing was corrected using the Benjamini-Hochberg method on the meta p-values and genes with an FDR < 0.05 were defined as cell type-specific marker genes. Cell types were identified based on cell type-specific marker genes from ImmGen (7), LungMAP (8), and Mouse Cell Atlas (3). Additional integrative methods, canonical correlation analysis and mutual nearest neighbors (9, 10), were also used to confirm and optimize cell clustering.

### Bulk RNA-seq

The UCLA Technology Center for Genomics & Bioinformatics (TCGB) generated libraries from total RNA and sequenced the libraries as paired-end 50 base pair reads using Hiseq 3000 (Illumina). Reads were aligned to Rnor 6.0 genome using HISAT2 version 2.1.0 and transcripts were assembled and quantified using StringTie version v1.3.3b (11).

### Identification of differentially expressed genes (DEGs) and pathways

DEG analysis was performed using MAST (12) and a false discovery rate (FDR) < 0.05 was considered statistically significant. Pathway enrichment analysis was performed using Gene Ontology (GO) (13), Hallmark pathways from Molecular Signature Database (14), Human Biomolecular Atlas Program (HuBMAP) (15), and DisGeNET (16) in R package fgsea v1.18.0 with genes ordered by MAST-derived Z scores. Human PAH-associated gene sets were obtained from DisGeNET (16) and Comparative Toxicogenomics Database (17) using the Harmonizome portal (18). For bulk RNA-seq data, DEG analysis was performed using DESeq2 (19) and a false discovery rate (FDR) < 0.05 was considered statistically significant. Enrichment of Hallmark pathways for overlapping genes between DEGs in EA2 (vs. EA1) and DEGs in human PAECs after either *BMPR2* knockdown or PMA stimulation were determined by hypergeometric test as implemented by Molecular Signatures Database at https://www.gsea-msigdb.org/gsea/msigdb/annotate.jsp.

### Independent datasets for validation

Select genes were queried in independent bulk and single-cell transcriptomic datasets of human lung and blood samples from Gene Expression Omnibus (GEO) or hosted on lab web servers. For the whole blood RNA-seq dataset (20), *Tm4sf1* expression was queried by functional class at https://sheffield-university.shinyapps.io/ipah-rnaseq-app/ to generate a PDF file showing individual datapoints along a log10(TPM) expression y-axis. The PDF was then imported into Inkscape version 1.1.1 after which the expression y-axis was proportioned to Inkscape y coordinates to determine expression values of individual data points.

---

**Table E1:**
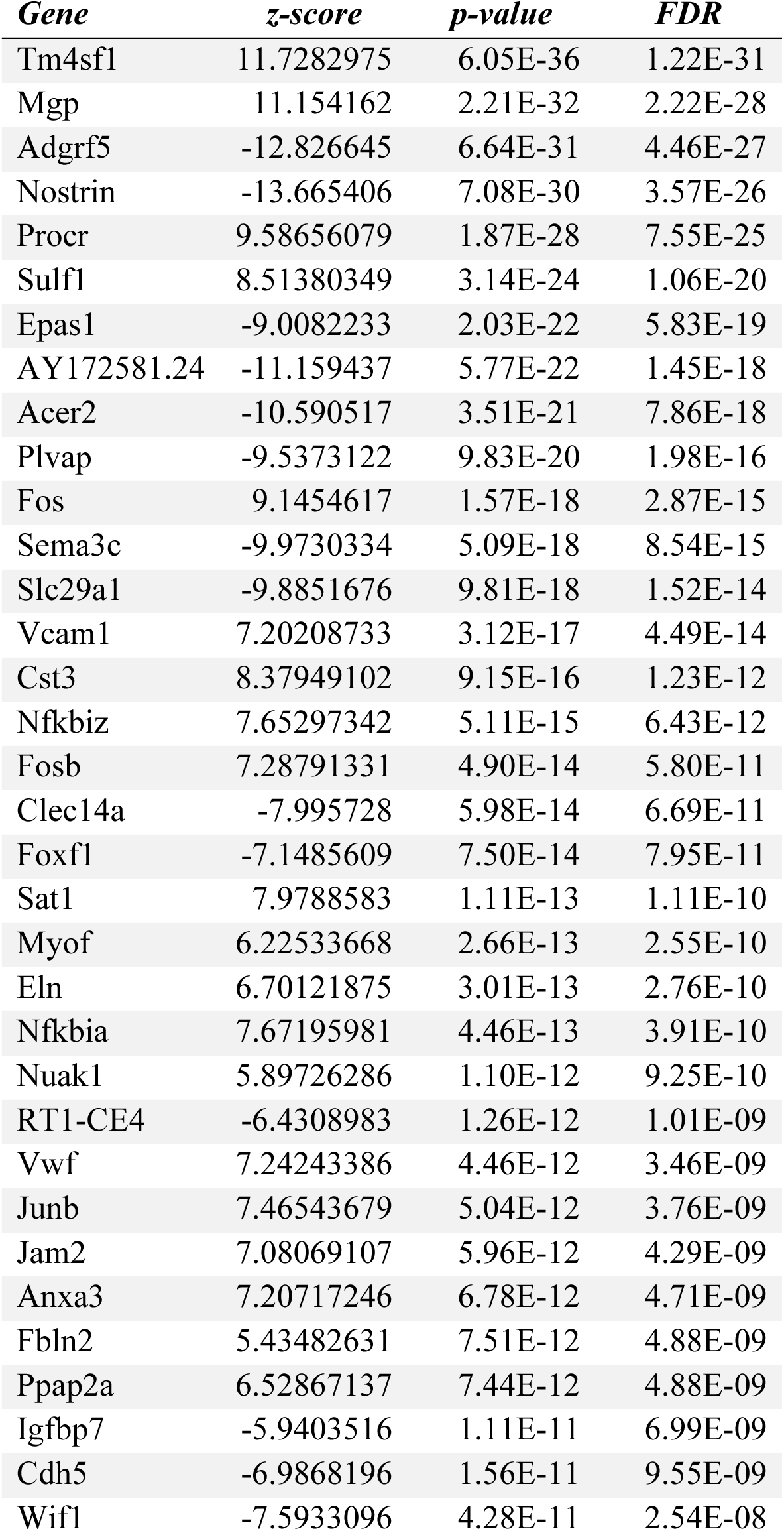
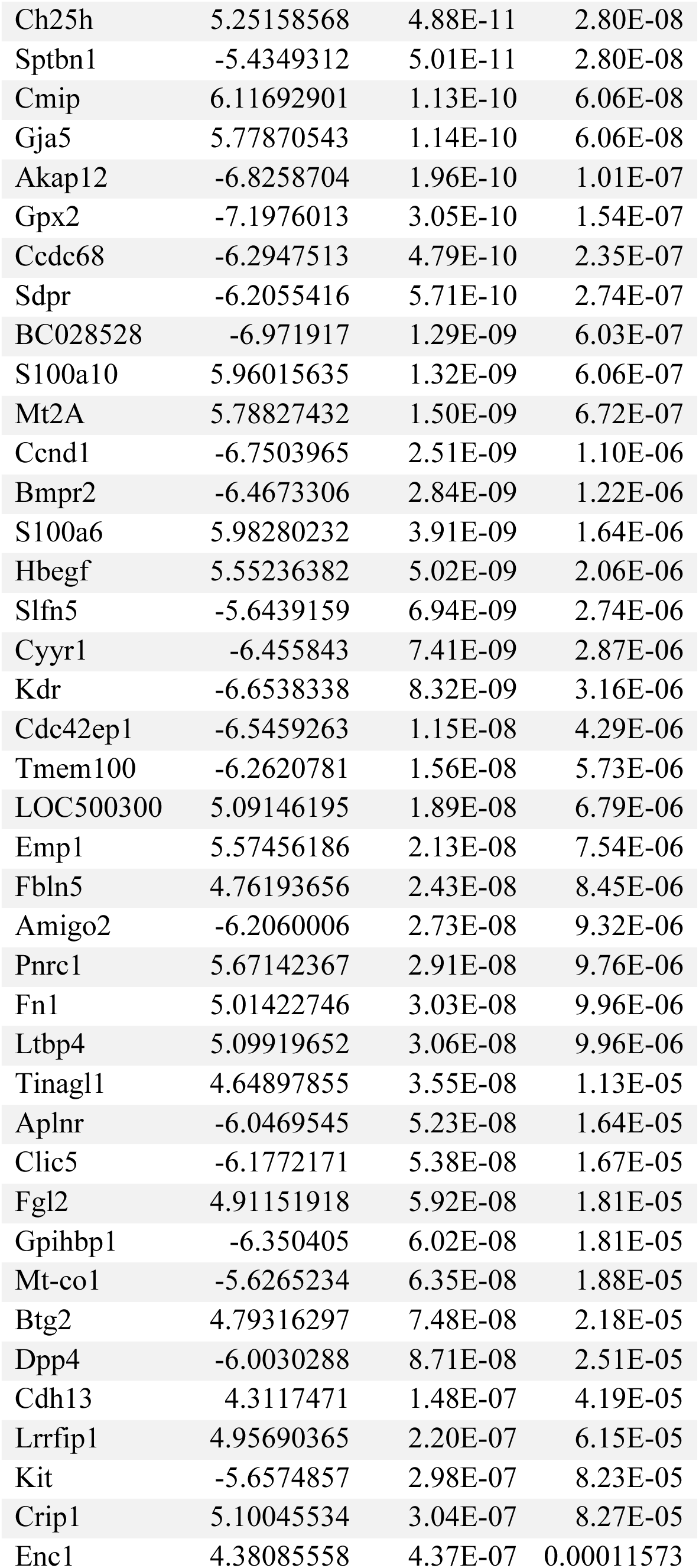
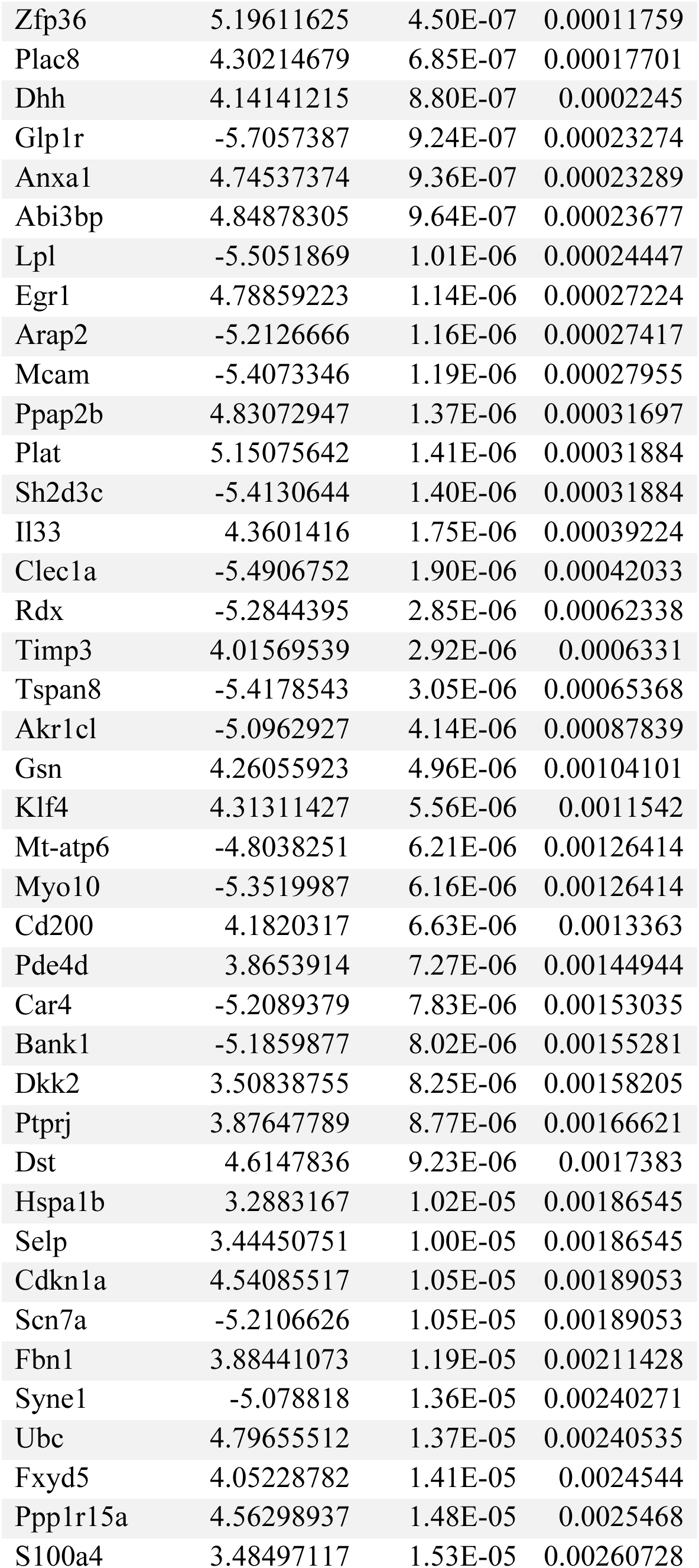
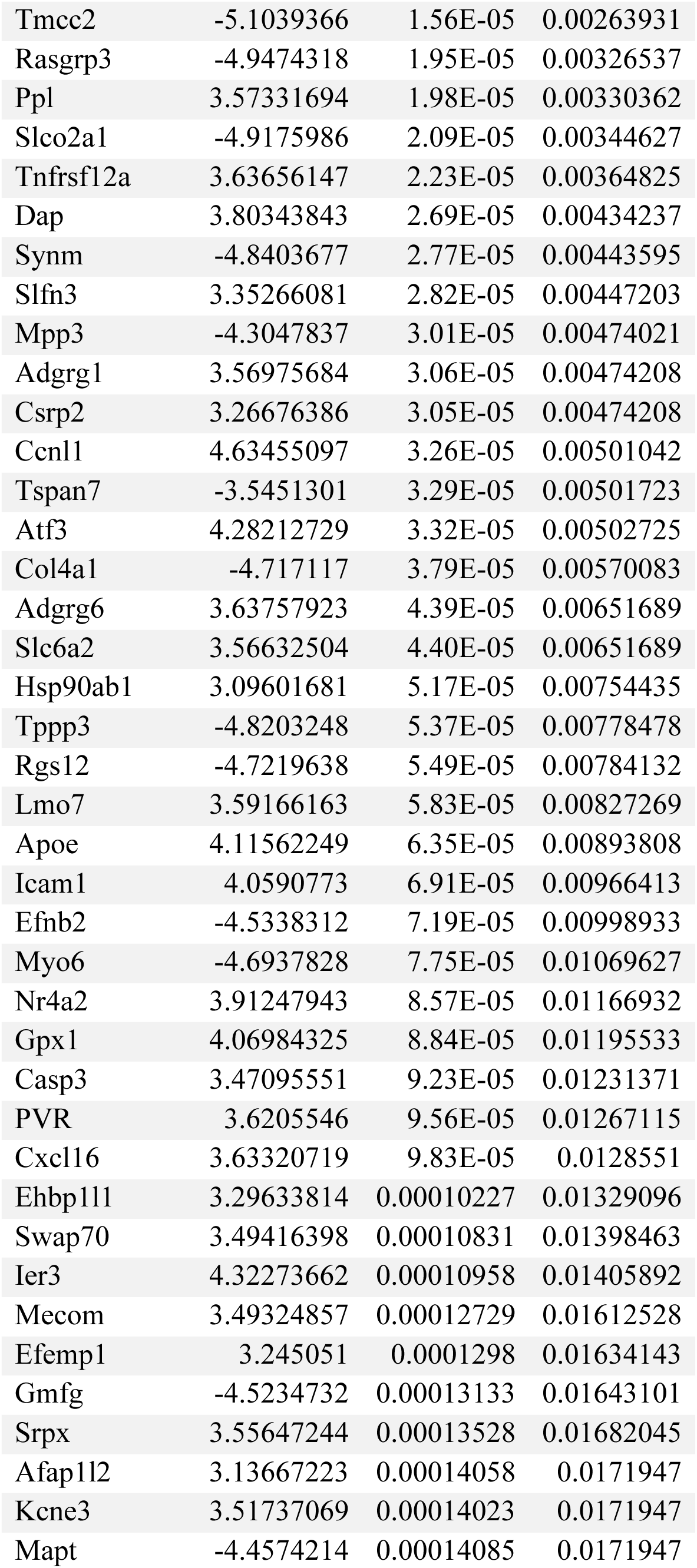
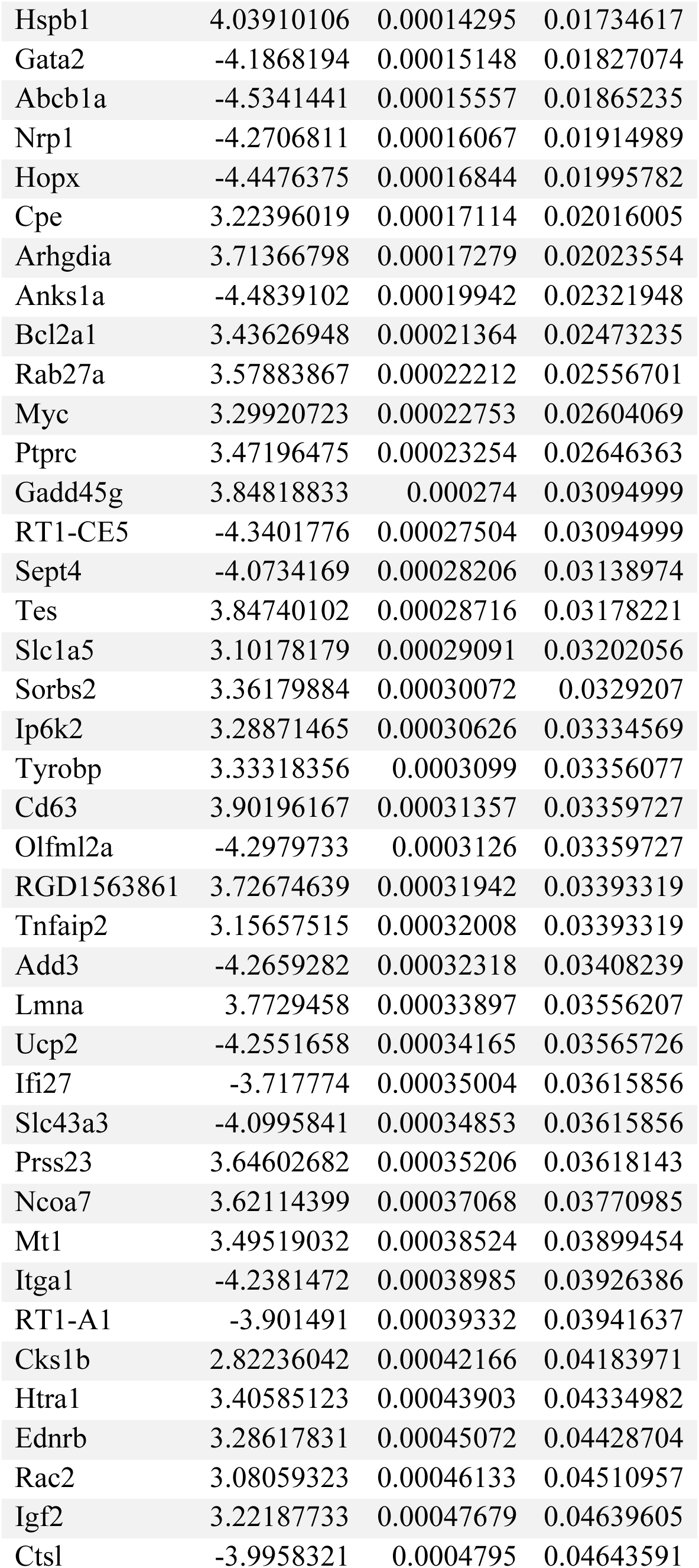
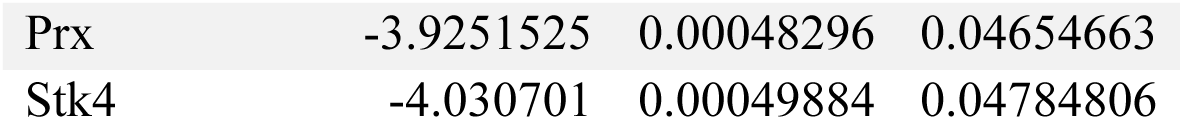
Genes differentially expressed in endothelial arterial type 2 (EA2) compared to endothelial arterial type 1 (EA1). Z-scores were derived from MAST where scores above 0 denote upregulated genes and scores below 0 represent downregulated genes. Genes with FDR < 0.05 are listed.

## Supplementary Figure Legends

**Figure E1:**
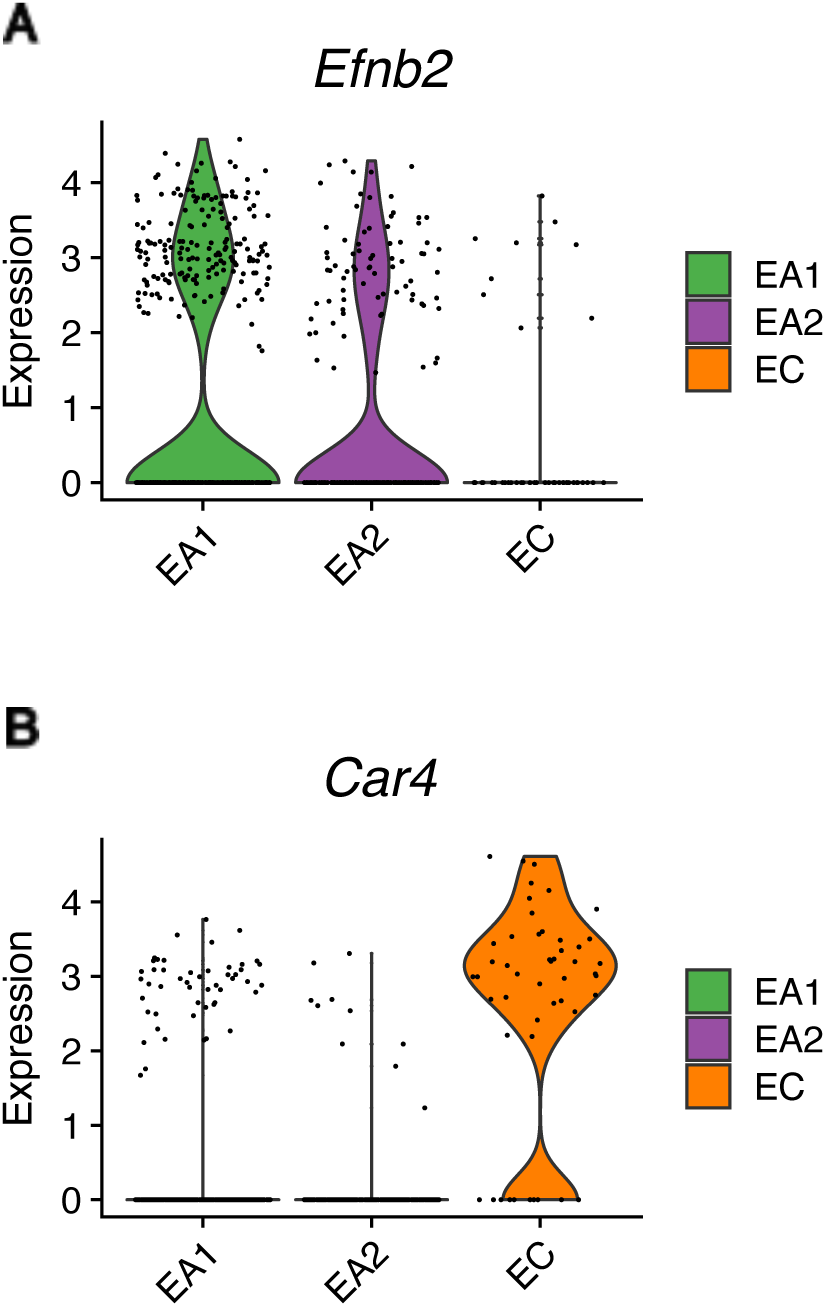
Endothelial expression of arterial marker *Efnb2* and capillary marker *Car4*. Violin plots showing expression of (A) *Efnb2* and (B) *Car4* in rat lung scRNA-seq grouped by endothelial subpopulations.

**Figure E2:**
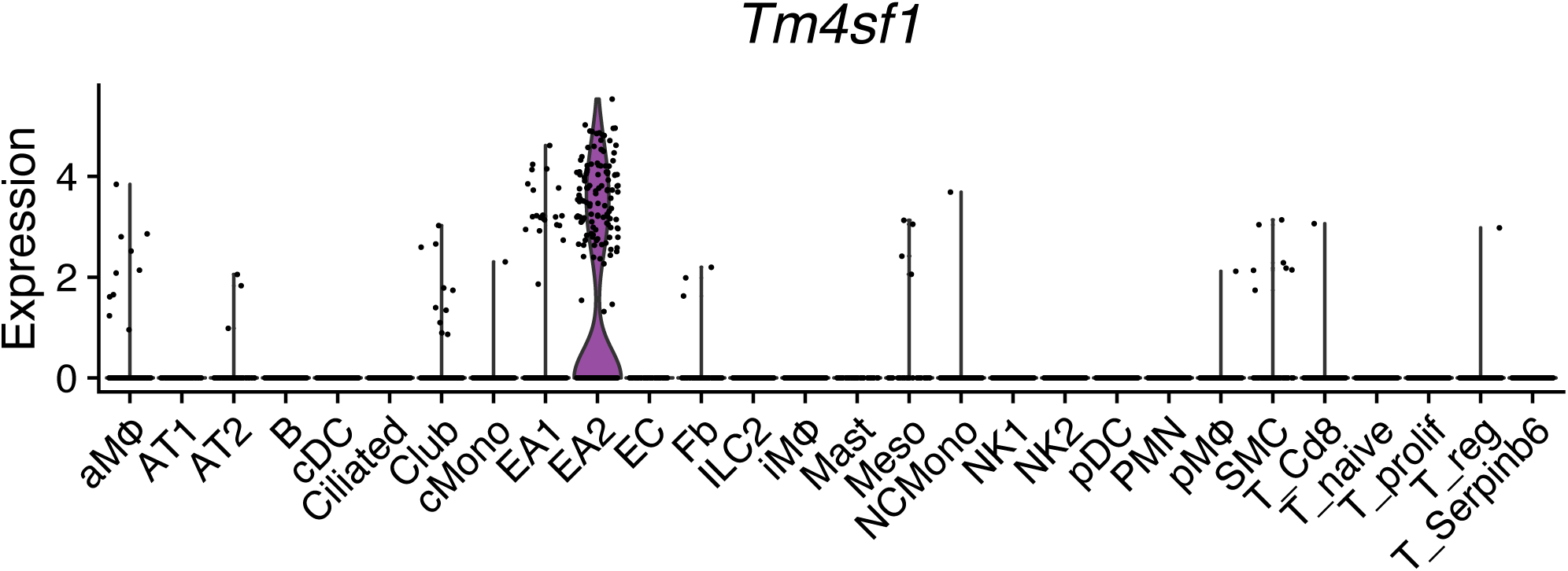
Violin plot showing expression of *Tm4sf1* across all rat lung scRNA-seq cell types.

**Figure E3:**
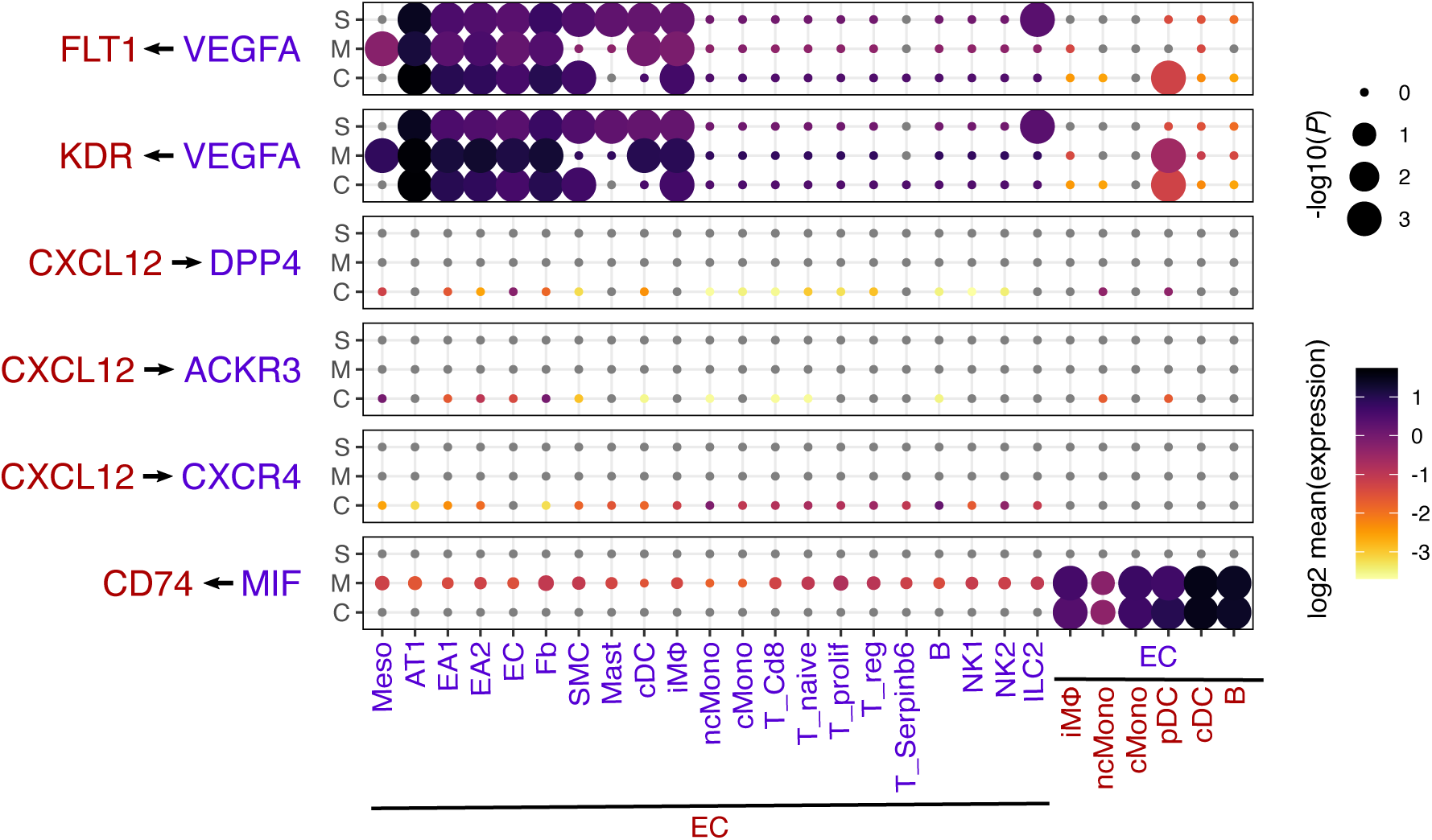
Ligand-receptor interactions in EC. Dot plot showing the log2 mean expression of select ligand-receptor pairs in control (C), MCT (M), and SuHx (S) between select cell types and EC. Size of dots are proportional to strength of p-value.

**Figure E4:**
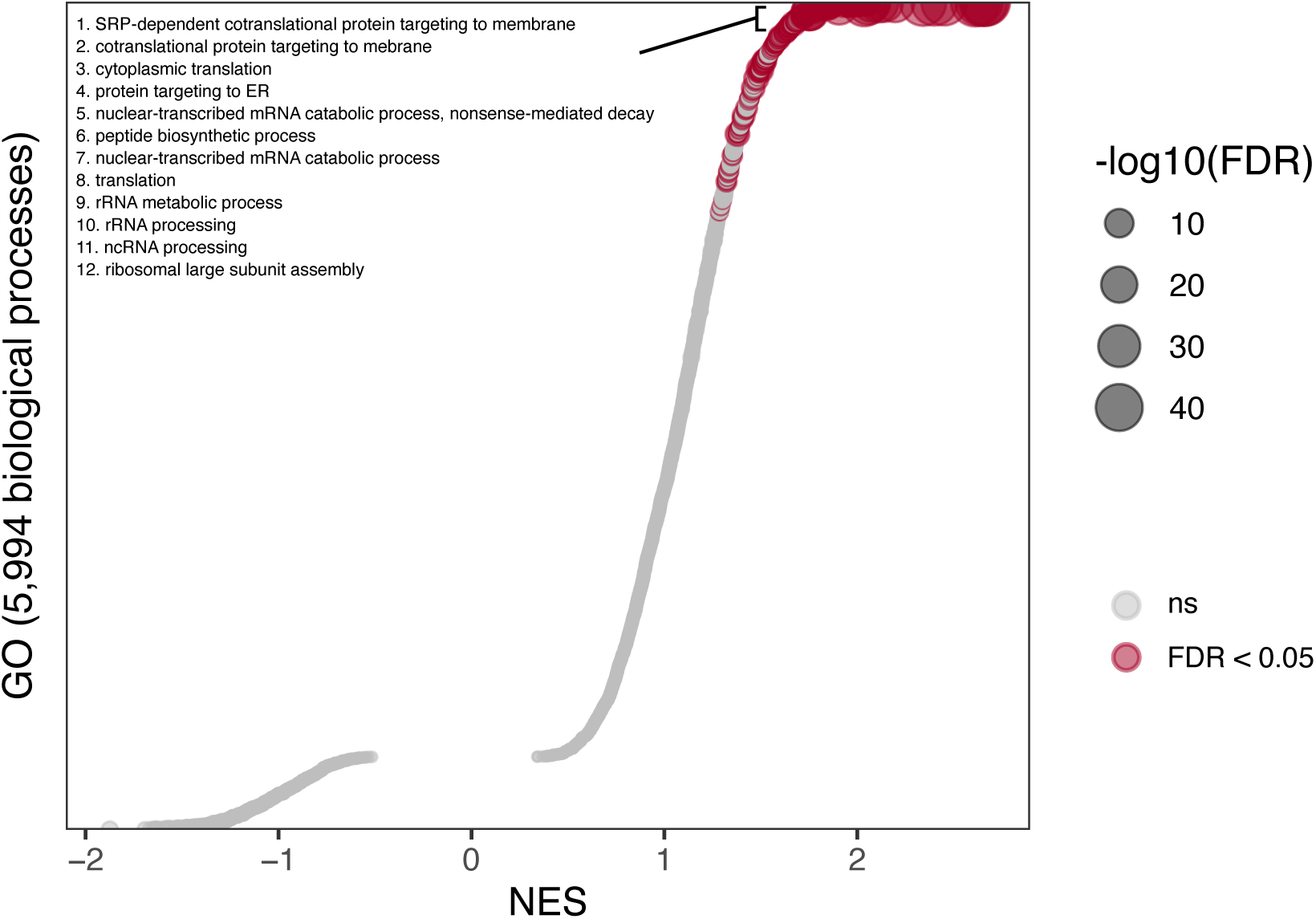
Biological processes enriched in genes correlated with a less differentiated state of endothelial cells. Dots plot showing GSEA of the CytoTRACE gene signature (i.e. genes correlated with CytoTRACE scores) using GO. The x-axis represents normalized enrichment scores (NES). Dots with FDR < 0.05 are colored in red and represent gene sets significantly enriched in genes correlated with differentiation state. The y-axis represents gene sets ordered by their NES. Select gene sets are labeled and numbered by their ordering as top gene sets enriched in genes correlated with a less differentiated state. Dots larger in size represent higher - log10(*FDR*) values.

**Figure E5:**
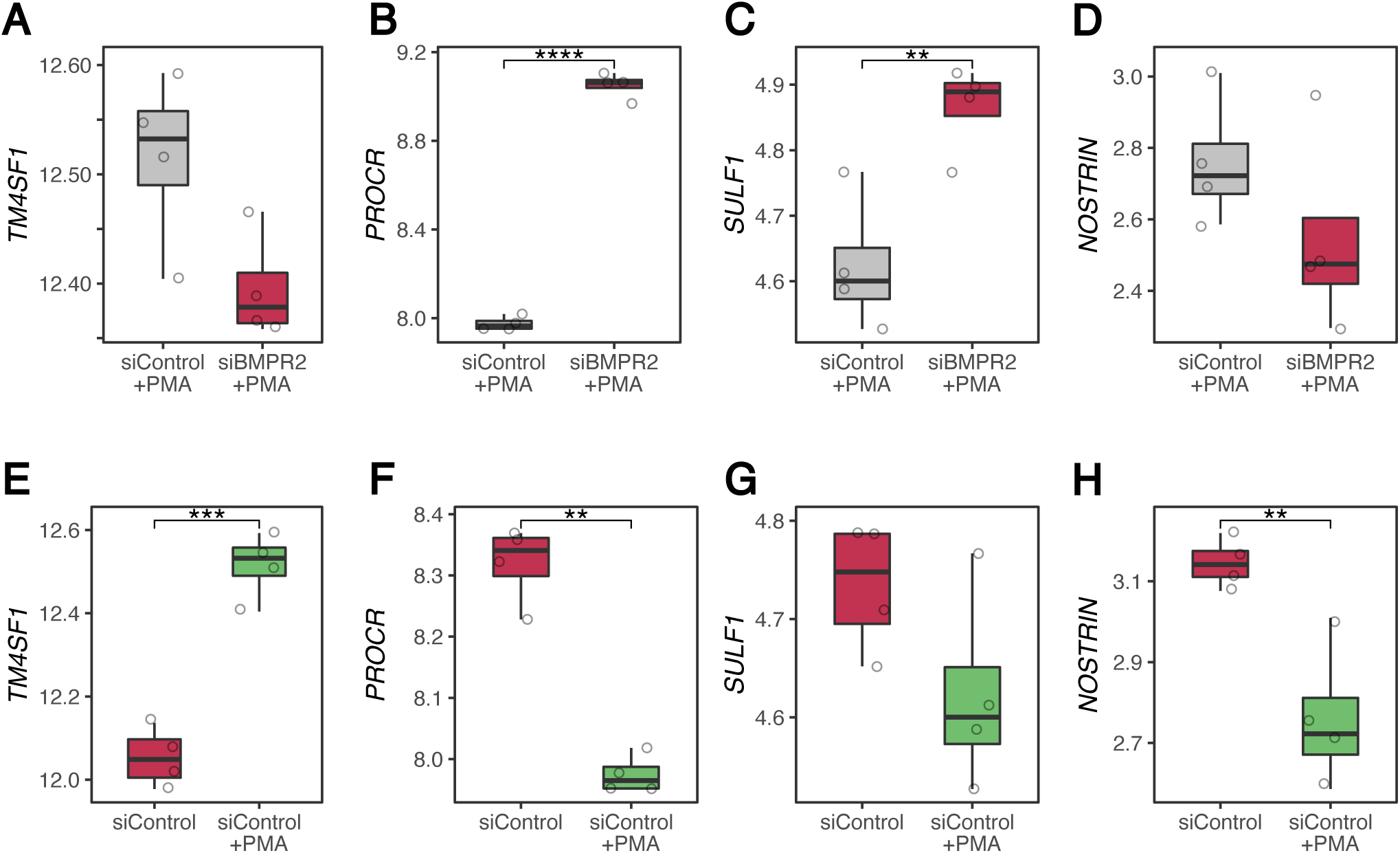
EA2 signature genes are differentially regulated by *BMPR2* knockdown and by PMA exposure in human endothelial cells. (A-D) Box plots showing expression of (A) *TM4SF1*, (B) *PROCR*, (C) *SULF1*, (D) *NOSTRIN* in 4 *BMPR2*- vs 4 control-siRNA-transfected primary human PAECs exposed to PMA from 4 donors (52). (E-H) Box plots showing expression of (E) *TM4SF1*, (F) *PROCR*, (G) *SULF1*, (H) *NOSTRIN* in 4 *PMA*-exposed vs 4 nonexposed siControl PAECs from 4 donors (52). PMA = phorbol 12-myristate 13-acetate.

**Figure E5:**
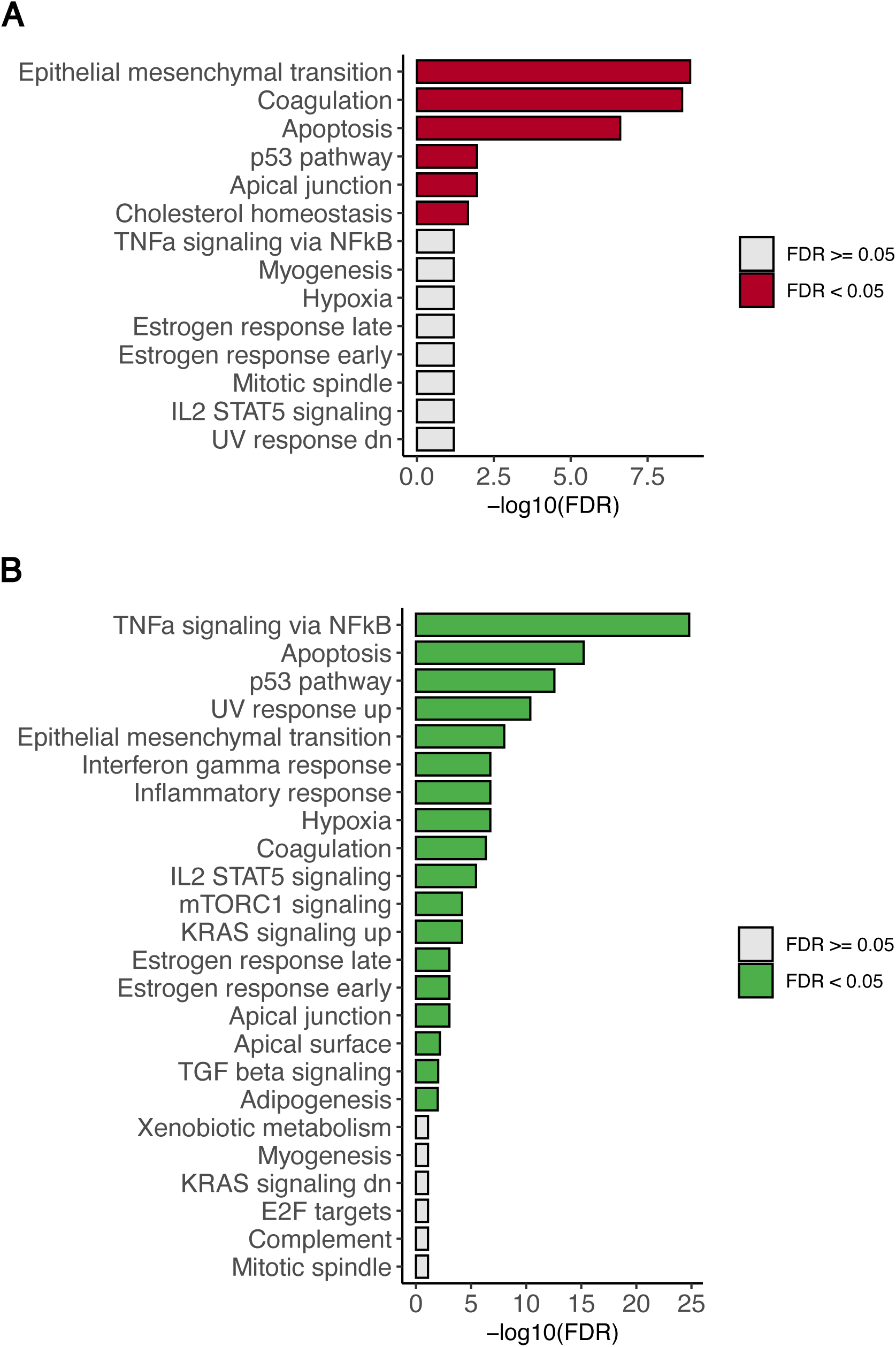
Overlapping DEGs between EA2 and *in vitro* PAH models are enriched for known PAH pathways. Bar plots showing enrichment of Hallmark pathways for overlapping genes between DEGs in EA2 (vs. EA1) and DEGs in human PAECs after either (A) *BMPR2* knockdown or (B) PMA stimulation. Enrichment was determined by hypergeometric test. FDR = false discovery rate.

## Notes

### Competing Interest Statement

The authors have declared no competing interest.

